# Inferring individual-level processes from population-level patterns in cultural evolution

**DOI:** 10.1101/111575

**Authors:** Anne Kandler, Bryan Wilder, Laura Fortunato

**Affiliations:** Department of Human Behavior, Ecology and Culture, Max Planck Institute for Evolutionary Anthropology; School of Engineering, University of Southern California; Institute of Cognitive and Evolutionary Anthropology, University of Oxford

**Keywords:** human cultural evolution, cultural micro- and macro-evolution, modes of cultural transmission, vertical, horizontal, oblique, mixed transmission, conformist bias, cultural diversity

## Abstract

Our species is characterized by a great degree of cultural variation, both within and between populations. Understanding how group-level patterns of culture emerge from individual-level behaviour is a long-standing question in the biological and social sciences. We develop a simulation model capturing demographic and cultural dynamics relevant to human cultural evolution, focusing on the interface between population-level patterns and individual-level processes. The model tracks the distribution of variants of cultural traits across individuals in a population over time, conditioned on different pathways for the transmission of information between individuals. From these data we obtain theoretical expectations for a range of statistics commonly used to capture population-level characteristics (e.g. the degree of cultural diversity). Consistent with previous theoretical work, our results show that the patterns observed at the level of groups are rooted in the interplay between the transmission pathways and the age structure of the population. We also explore whether, and under what conditions, the different pathways can be distinguished based on their group-level signatures, in an effort to establish theoretical limits to inference. Our results show that the temporal dynamic of cultural change over time retains a stronger signature than the cultural composition of the population at a specific point in time. Overall, the results suggest a shift in focus from identifying the one individual-level process that likely produced the observed data to excluding those that likely did not. We conclude by discussing the implications for empirical studies of human cultural evolution.

## 1 Introduction

Our species is characterized by a great degree of cultural variation, both within and between populations. This variation can be observed in domains as different as material culture [e.g. 1], linguistic features [e.g. 2], or social norms [e.g. 3]. These population-level patterns are the aggregate product of underlying individual strategies. Understanding how those patterns of culture emerge from individual-level behaviour is a longstanding question in the biological and social sciences [4].

The field of cultural evolution encompasses efforts to answer this question, through a variety of theoretical and empirical tools [see 5, for a recent review]. Cultural evolution is the process of change in the frequency of different variants of a cultural trait over time. Cultural traits comprise the knowledge, ideas, beliefs, skills, attitudes, or any other form of information that can be socially transmitted between individuals, for example through teaching or imitation [6]. Seminal early contributions to the field focused on human culture [e.g. 7,8], but the scope now extends to non-human culture as well [9].

Overall, this body of work has identified a number of factors shaping cultural variation within and across human groups, including the pathways for the transmission of information between individuals, demography, shared population history, and adaptation to environmental conditions. However, we lack a systematic understanding of the effect of these different factors and their interactions. In particular, we lack a conceptual framework explicitly mapping the patterns observed at the level of groups onto the underlying processes occurring at the level of individuals [5]. For example, a large body of empirical work in cultural evolution builds on the notion of the relative “conservativeness” of vertical transmission (i.e. parent to child) compared to other transmission pathways (e.g. horiziontal transmission, i.e. between peers) [7]. This body of work includes field-based investigations [e.g. 10, 11, 12] and cross-cultural studies [e.g. 13, 14, 15, 12]. But how exactly does the group-level “signature” of vertical transmission differ from that of other transmission pathways? And is it sufficiently different that it can be mapped, unequivocally, onto vertical transmission?

To address these and related questions, we develop a simulation model capturing demographic and cultural dynamics relevant to human cultural evolution, focusing on the interface between population-level patterns and individual-level processes [see e.g. 16, 17, 18, 19, for other modelling frameworks]. By design, the simulation model is the simplest it can be. For example, we focus on neutral cultural traits (i.e. not linked to to fitness) and constant population sizes; traits only differ in the way they are transmitted between individuals. In this respect, our work differs from related recent analyses by [18], which explored instead the effect of trait transmission on the age-structure of a population, and the extent to which cultural traits affecting demographic change can spread.

In other words, our model does does not seek to replicate a specific cultural system. Rather, it is used to run “artificial experiments” for various demographic and cultural scenarios, in order to explore and analyse the ranges of possible evolutionary outcomes generated by those scenarios. Specifically, the model tracks the distribution of variants of cultural traits across a population over time, conditioned on different pathways for the transmission of information between individuals. From these simulated data we obtain theoretical expectations, in the form of probability distributions, for a range of statistics commonly used in the literature to capture population-level characteristics (e.g. the degree of cultural diversity).

Our aim is two-fold. First, we aim to derive general insights about the different transmission pathways and their signatures at the level of the group. Second, we aim to establish whether, and under what conditions, the different pathways can be distinguished based on their group-level signatures. Are the signatures sufficiently different that they can be traced back to specific individual-level processes? The rationale is that if we cannot accurately infer underlying scenarios from simulated data, which are generated under known “experimental” conditions, it is unlikely that we will be able to do so based on empirical data.

Some additional context may help grasp the significance of the second aim. Researchers in anthropology and related disciplines often draw inferences from data on variation in cultural traits within a population, but these data are typically sparse in space and/or time. Our analysis aims to establish the theoretical limits to inference — in other words, how much information can in fact be “extracted” from the data under simplified scenarios of cultural change captured by our simulation framework. These theoretical limits represent an upper-bound to inference: we cannot expect to obtain more information from empirical data, which are the product of intricate real-world scenarios.

We begin by describing the mathematical framework and the statistical analysis (Section 2), followed by presentation of the results (Section 3). The code used to generate the results is available from https://github.com/bwilder0/cultural-evolution. We conclude with a non-technical summary of the key findings, together with discussion of their implications (Section 4); readers can skip to this part without loss of insight.

## 2 Methods

Our analysis involves two components. The first is a mathematical framework simulating human cultural evolution. The second is a set of statistics used to summarize the simulation output; these are used to determine whether the group-level patterns extracted from the simulation output can be reliably distinguished.

The framework includes both discrete and continuous cultural traits: discrete traits assume a finite number of variants, continuous traits assume any value in a given interval. For ease of presentation, this and the next section focus on the discrete case. An outline of key results for the continuous case are in Section 3.2.4, with additional detail in Section S3.3.4 of the supplementary material.

### 2.1 Mathematical framework

We consider a simple age-structured population, divided into five age classes, where each individual possesses a variant of five different cultural traits.

#### 2.1.1 Cultural traits and transmission modes

Discrete cultural traits assume one of five possible variants. As the traits are assumed to be neutral, individuals do not benefit from carrying a specific variant over another.

Following [7], we define a set of modes for the transmission of trait variants between individuals (Table 1). Broadly, the modes describe the flow of information within and between generations. They delimit the set of potential interaction partners within a population, with interactions between individuals within this set occurring at random.

**Table 1:**
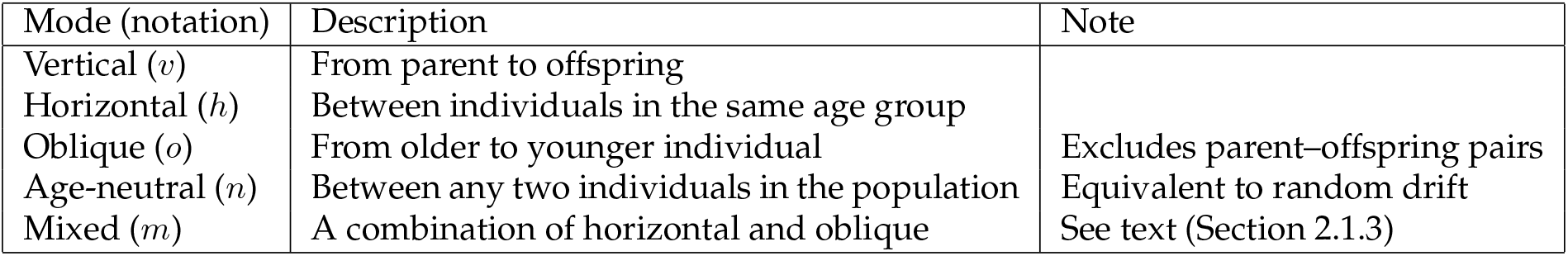
Transmission modes.

Additionally, transmission biases may steer interactions towards individuals carrying a particular variant [8]. A large number of biases have been identified in the literature [see 20]; we focus on conformity bias, i.e. a preference for the most common variant in the population, as detailed in Section 2.1.4.

We consider a population of *N* individuals. At each point in time individual *i* is described by a variable 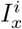, with 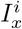 ∈ {1,…, 5} for *x = v,h, o, n, m* (Table 1). This is the individual’s current cultural make-up, comprising five cultural traits, each linked to a transmission mode. For all traits, an individual acquires the variant of its parent at birth, i.e. through vertical transmission. For all traits except the one linked to mode *v*, the variant may change through further transmission events during the individual’s lifetime, as described in Section 2.1.3.

#### 2.1.2 Demographic processes

In addition to variable 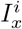, at each point in time individual *i* is described by a variable 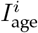, denoting its current age group. There are five age groups, covering the age range 0-50. The age structure of the population is summarised by [*n*_1_*,…, n*_5_], where *n_i_* describes the number of individuals in age group *i*. It holds 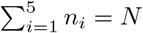.

Each time step in the simulation corresponds to 10 years. At each time step, individuals age by moving to the next age group, and they may either die or reproduce. We assume a constant population size *N*. Therefore, the number of individuals entering the population through reproduction in a given time step is determined by the difference between *N* and the number of deaths in the previous time step. Individuals die with probability *p*_death_. All individuals in age group 5 die with *p*_death_ = 1 in the following time step.

Reproduction occurs asexually. Individuals in age groups 2 and 3 are chosen at random at each time step to reproduce until population size *N* is reached. These individuals produce offspring who inherit their trait variants. However, with a small probability *μ* a mutation occurs and the offspring’s variant differs from the parent’s, as described in Section 2.1.3.

These demographic processes produce a pyramid-shaped age structure, with fewer individuals in older generations. It is possible that all potential parents die before reproducing, leading to extinction of the population. The results presented below are based on simulation runs where the population survived for the entire duration of the simulation.

#### 2.1.3 Mutation and transmission events

We define a mutation rate *μ*, which describes the fidelity of the transmission process and applies to all transmission events. For all traits, individual *i* acquires the variant of its parent with probability 1 – *μ*, and a randomly selected variant with probability *μ*. This process describes vertical transmission (Table 1).

Additionally, for traits linked to modes *h, o, n* the variant may change through further transmission events during the individual’s lifetime. Specifically, at each time step individual *i* engages with probability *p*_w_ in interaction with a randomly selected individual *j*. For example, *p_w_ =* 0.5 corresponds to an average of 2.5 transmission events during the individual’s lifetime, *p*_w_ = 1 to an average of 5 events. During each transmission event, individual *i* acquires the variant of individual *j* with probability 1 – *μ*, and a randomly selected variant with probability *μ*. The transmission rules in Table 2 apply to these interaction events; they show that horizontal, oblique, and age-neutral transmission differ only in the set of potential interaction partners (Section 2.1.1).

**Table 2:** Transmission rules for modes *h, o, n* in the absence of mutation.

| Mode | Possible interaction partner $j$ for individual $i$ | Transmission rule |
| --- | --- | --- |
| $h$ | $I_{\text{age}}^i = I_{\text{age}}^j$ | $I_h^i = I_h^j$ |
| $o$ | $I_{\text{age}}^i > I_{\text{age}}^j$ | $I_o^i = I_o^j$ |
| $n$ | any individual | $I_n^i = I_n^j$ |

The “pure” modes *v, h, o, n* provide a useful baseline against which to compare more realistic scenarios. Specifically, one of the five traits in an individual’s cultural make-up is linked to mode *m*, whereby possible interaction partners include both peers and older individuals (Table 1). We term this mode “mixed”, to indicate that it is effectively the combination of horizontal and oblique transmission. For this trait, individual *i* engages in horizontal transmission with probability (1 – *p*_mix_) and in oblique transmission with probability *p*_mix_, following the rules specified in Table 2. Qualitatively, the higher the value of *p*_mix_, the more likely that transmission is oblique, i.e. that the individual’s interaction partner belongs to an older age group, rather than to the same age group.

Across all modes, the interaction probability *p*_w_ and the mutation rate *μ*, combined, determine the potential for cultural change. Broadly, we distinguish between scenarios with low vs. high potential for cultura change (e.g. low *μ*, low *p*_w_ *vs.* high *μ*, high *p*_w_). For instance, the higher *p*_w_ and *μ*, the more opportunities there are for an individual’s cultural make-up to change over its lifetime, and hence for the cultura composition of the population to change with each time step.

#### 2.1.4 Conformity bias

As noted in Section 2.1.1, the transmission modes describe different sets of potential interaction partners but interactions between individuals within each set occur at random. This implies that the probability *p_k_* that an individual interacts with a partner carrying variant *k* is proportional to the relative frequency of variant *k* in the set. It holds

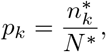

where *n*_k_* describes the number of individuals in the set of size *N\** carrying variant *k*.

Conformist transmission is defined as the disproportional adoption of common variants [8]. With conformity bias, the probability of an individual interacting with a partner carrying variant *k* is given instead by [21, 22]

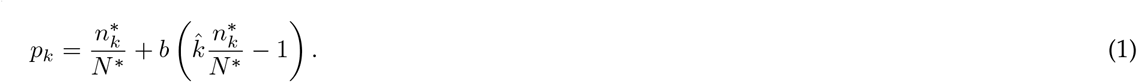

The parameter *b* ≥ 0 describes the strength of the bias and *k̂* is the number of variants present in the population. Conformity increases the probability that an individual carrying variant *k* acts as interaction partner if the frequency of variant *k* exceeds the “relative” majority 1 /*k*̂.

#### 2.1.5 Simulation set-up

At the beginning of each simulation run the *N* individuals in the population are distributed randomly across the five age groups. They all carry the same variants of the cultural traits. In each subsequent time step the demographic and cultural processes described above take place, over time leading to changes in the frequency of the different variants, and thus to the cultural composition of the population.

Individuals update their cultural make-ups asynchronously, i.e. an individual’s interaction partner may have already updated its cultural make-up in the given time step. The order in which individuals engage in transmission events is randomised in every time step.

A single simulation run consists of a burn-in phase followed by 200 time steps. We explore various constellations of the parameters for ranges *N* = 25, 50, 100; *μ* = 0.01, 0.05, 0.1; *p*_w_ = 0.5, 0.75, 1; *b* = 0, 0.01, 0.02, 0.03. Note that *b* = 0 for all analyses, except those investigating specifically the effect of conformity on distinguishability of the different transmission modes.

### 2.2 Statistical analysis

#### 2.2.1 Summary statistics

The mathematical framework tracks the frequencies of trait variants at each time step. We use this information to characterize the cultural composition of the population and the dynamic of cultural change ove time, conditioned on the different transmission modes. Specifically, with *x* = *v, h, o, n, m* denoting the mode (Table 1), for a given trait we derive

i. the frequencies of its five variants [*p_1_^x^, p_2_^x^, …, p_5_^x^*] across individuals in the population, where *p_k_^x^* describes the relative frequency of variant *k*,
ii. the frequency of the most common variant in the population, denoted 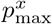,
iii. the total number of variants present in the population, denoted *k^x^*,
iv. the level of cultural diversity, as measured by the Shannon diversity index

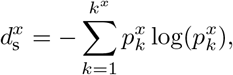

where *k^x^* describes the number of variants present and p_k_^x^ the relative frequency of variant *k*, as defined above,
v. the average time a variant stays the most common variant, denoted 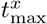.

The first statistic is the joint probability distribution of the frequencies of the five cultural variants. While this captures the most information about the cultural composition of the population at a given point in time, researchers may not have access to the full data. Accordingly, we also explore how much information can be extracted from “partial” data, namely the frequency of the most common variant, 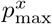, and the total number of variants present, *k^x^*.

Researchers often summarise the cultural composition of a population using a diversity measure such as the Shannon diversity index [e.g. 1, 23, 24]. We include this statistic in our analysis to investigate how the dimension reduction this involves affects our ability to distinguish between transmission modes.

Finally, we characterise the temporal dynamic of cultural change with 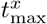, which measures the rate of change of the most common variant in the population. Broadly, this gives an indication of how fast the cultural composition of the population can change over time. We use 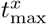 to explore whether the temporal dynamic is more informative than the cultural composition of the population at a given point in time, as described by the other statistics.

#### 2.2.2 Distinguishability analysis

We explore the behaviour and inferential power of each statistic by generating probability distributions of the statistic, across simulation runs, for different values of the parameters in the model.

For a given parameter constellation, we determine whether two modes can be distinguished based on the statistic by (i) examining the area of overlap between the corresponding distributions (e.g. vertical vs. horizontal transmission) and (ii) determining the probability that a particular transmission mode acted in the population to produce on observed value of the statistic. The approach can be applied more generally to determine the effect of a given parameter. This is done by comparing the distributions for a given mode under different values of the parameter (e.g. oblique transmission with *vs.* without conformity, or vertical transmission with small vs. large population size), keeping constant the values of the other parameters. For ease of presentation, we focus here on comparison of two modes for a given parameter constellation. For simplicity, we describe the procedure for one-dimensional probability distributions (i.e. the distributions for 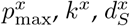 and 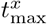), but the same approach applies to the joint distributions *f_joint, x_*.

We take the area of overlap between the distributions to indicate to what degree the two corresponding modes can be distinguished based on the statistic without knowledge of an empirical estimate. At one extreme, no overlap suggests that the modes can be reliably distinguished; at the other, complete overlap suggests that they cannot be distinguished.

In detail, the area of overlap *O_xy_* between probability distributions *f_x_*and *f_y_* is determined using the Kolmogorov-Smirnov distance *K_xy_* between the associated distribution functions *F_x_* and *F_y_* (Figure 1(a)). If *z*\* denotes the value where 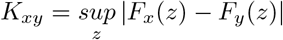 is realized, then it holds [25]

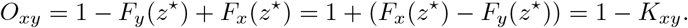

**Figure 1:**
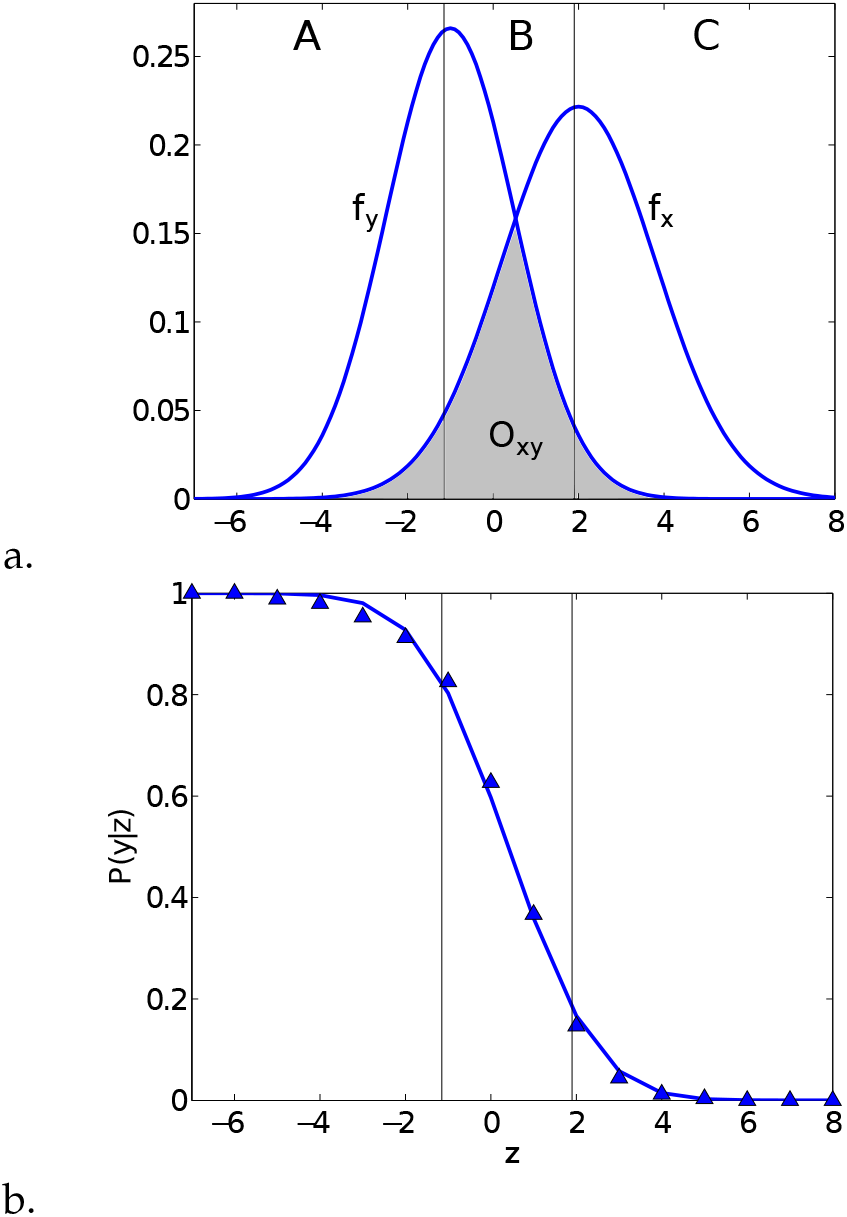
Illustration of (a) the area of overlap *O*_*xy*_ between probability distributions *f*_*x*_ and *f_y_*, and (b) the probability *P*(*y*|z) that transmission mode *y* (as opposed to *x*) acted in the population to generate value *z*.

Distinguishability is defined using a threshold value *Ō*: two distributions *f*_*x*_ and *f*_*y*_ are distinguishable if it holds *O_xy_* < *Ō*, and we use a value of *Ō* = 20%, corresponding to the widely accepted 80% power cut-off. An alternative approach, based on the Rayleigh criterion, is introduced in Section S1 in the supplementary material.

Specifically, based on 30,000 simulation runs we determine (i) the joint distributions f _joint,*x*_ of frequencies of the five trait variants, and (ii) the probability distributions 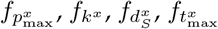 of statistics 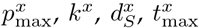, conditioned on transmission modes *x* = *v,h, o, n, m*. For a given statistic we calculate the areas of overlap *O*_*xy*_ for pairs of distributions *x* and *y*, and we compare their values against the threshold *Ō* = 20% to determine whether the corresponding modes are distinguishable based on the statistic.

This procedure rests on an *a priori* definition of distinguishability, without considering available empirical data. The results provide general expectations about the similarity or dissimilarity of population-level outcomes generated by the different transmission modes. Even in the absence of empirical data, these expectations can inform researchers as to whether the corresponding statistics 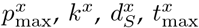 carry a signature of the underlying transmission modes.

If researchers do have access to empirical estimates of a statistic, then the procedure can be extended to incorporate this information. This is illustrated with an example in Figure 1(a). Values of the statistic in regions A and C are almost “unique” to modes *x* and *y*, respectively, whereas values in region B are “shared”. The interpretation is that values of the statistic in regions A and C could only have been produced by one of the two modes, whereas values in region B could have been produced by either mode. It follows that empirical estimates provide no additional information in cases where the area of overlap *O_xy_* is close to the extremes 0 and 1 (corresponding to no overlap and complete overlap of the distributions, respectively).

In detail, given the empirical estimate of a statistic, denoted by *z*, the aim is to determine the probability that a given transmission mode acted in the population to produce this value. We define the set of possibletransmission modes Γ = {*v, h, o, n, m*} and assume that one of these did produce the observed value of the statistic. Bayes′ theorem then results in

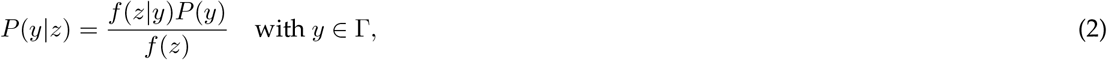

where *P*(*y* |*z*) stands for the probability that mode y is acting in the population given *z* (see blue triangles in Figure 1(b)). The probability distribution *f* (*z*|*y*) is the distribution generated by the simulation framework; it captures the values of the statistic that can be assumed for mode *x*. The probability *P* (*y*) includes all prior information about the likelihood that mode *y* acted in the population; in the absence of prior information we assume that 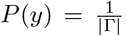. The function *f* (⋅) describes the probability distribution of the values of the statistic and it holds 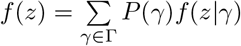.

For comparison of two transmission modes, the probability *P*(*y*|*z*) can be determined by logistic regression (see blue line in Figure 1(b)). Multinomial logistic regression can be used for more than two modes. Probabilities close to 0 or 1 have a clear interpretation: at 0, mode *y* could not have produced the observed value of the statistic; at 1, mode *y* is most likely to have produced the observed value, compared to the possible alternatives. Intermediate values indicate that multiple transmission modes could have produced the observed value of the statistic. Alternatively, receiver operating characteristic (ROC) curve analysis could be applied (see Section S2 in the supplementary material for details).

## 3 Results

### 3.1 Population-level patterns

In this section we explore whether the different transmission modes result in characteristic population-level outcomes under scenarios with low vs. high potential for cultural change, as defined in Section 2.1.3. An example of the analyses underlying the results is presented in Section S3.1 in the supplementary material.

#### 3.1.1 Pure modes

We begin by investigating the behaviour of the pure modes — vertical, horizontal, oblique, and age-neutral transmission. To this end, we study the distributions of statistics 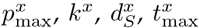 conditioned on transmission modes *x* = *v, h, o, n*, for parameter constellations *N* = 25, 50, 100; μ = 0.01, 0.05, 0.1; *p*_w_ = 0.5, 0.75, 1.

Statistics 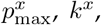 and 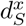 describe the cultural composition of the population at a given point in time (Section 2.2.1). The different transmission modes result in comparable population-level outcomes for these statistics under scenarios with low potential for cultural change. At the same time, outcomes for oblique transmission are strongly affected by the value of *p*_w_. Specifically, under scenarios with high potential for cultural change oblique transmission leads to more homogenous cultural compositions than vertical, horizontal, and age-neutral transmission.

These differences in behaviour across transmission modes are rooted in the different sets of potential interaction partners. In age-neutral transmission an individual can interact with any other individual in the population. In horizontal transmission an individual in age group k can only interact with the *n_k_* – 1 individuals in its own age group (corresponding to age-neutral transmission within the age group). In oblique transmission an individual in age group *k* can interact with 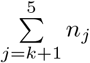 older individuals, where *n_j_* is the number of individuals in age group *j* (corresponding to age-neutral transmission within all age groups older than the individual’s). In other words, compared to age-neutral and horizontal transmission, oblique transmission is characterised by a disproportional influence of older individuals. Individuals in age group 5 are potential interaction partners for all individuals in age group 4 and younger, whereas individuals in age group 2 are potential interaction partners only for individuals in age group 1. As a result, a variant at high frequency in age group 5 tends to “trickle down” to the other age groups; over time, it tends to become increasingly common, eventually leading towards homogenisation of the population as a whole.

To illustrate the influence of the age structure on the transmission dynamic we derive the probability that a mutant variant (i.e. a variant with frequency 1) has frequency 0, 1, 2, …, *N* after one time step (Section S3.2 of the supplementary materials). For oblique transmission the spread probability of a mutant variant is highly influenced by the age of the individual introducing it into the population: the older the individual, the higher the probability that the variant is still present in the population after one time step. By contrast, for horizontal transmission the spread probability does not vary greatly with the age of the individual introducing it into the population. By definition, the same is true for age-neutral transmission.

How does this translate into population-level outcomes? Figure 2 shows the level of cultural diversity 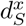 for the different modes in a scenario with high potential for cultural change (parameter constellation *N* = 100, *μ* = 0.1, *p_w_* = 1), separately in each of the five age groups and in the population as a whole. For oblique transmission the level of diversity within age groups is comparable to the level of diversity in the whole population. This suggests that the age groups are culturally more homogeneous, as expected based on the trickle-down effect. The same pattern applies to age-neutral transmission; by definition, this mode is not affected by the age structure of the population. By contrast, for horizontal transmission the level of diversity within age groups is substantially lower than the level of diversity in the whole population. This suggests that the different age groups sustain different cultural variants.

**Figure 2:**
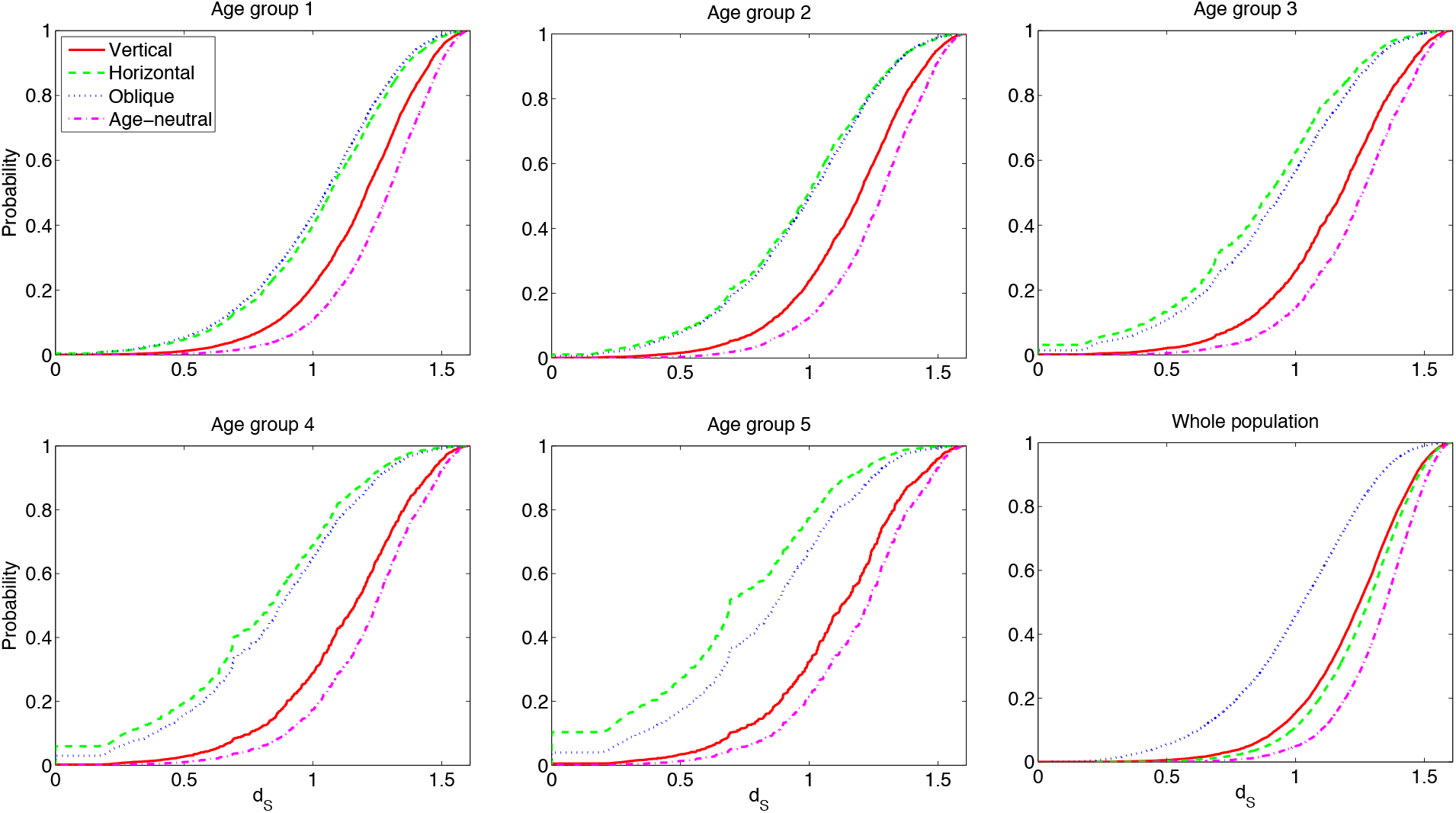
Level of cultural diversity in the five age groups and in the whole population for the pure transmission modes. Shown are the distribution functions of statistic 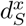 for the different modes, under parameter constellation *N* = 100, *μ* = 0.1, *p_w_ =* 1.

A final insight relates to the relative rate of cultural change for the different transmission modes. Specifically, the modes can be ranked by ordering the distribution functions of 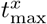 (Section S3.1.1 in the supplementary material). As expected, vertical transmission leads to the slowest rate of change, as there are limited opportunities for transmission compared to the other modes (see Figure S1 in the supplementary material). Further, oblique transmission is characterised by a slower rate of change than horizontal and age-neutral transmission, due the disproportional influence of older age groups and the consequent homogenisation of the population.

#### 3.1.2 Mixed mode

The results for mixed transmission reveal a consistent pattern (Section S3.1.2 in the supplementary material). At low values of *p*_mix_ the occasional oblique transmission event may introduce variants otherwise absent in a given age group. This leads to an increase in cultural diversity at the population level compared to pure horizontal transmission. At high values of *p*_mix_ the occasional horizontal transmission event effectively dampens the trickle-down effect, weakening the disproportional influence of older age groups and the consequent homogenization of the population. This leads to an increase in cultural diversity at the population level compared to pure oblique transmission. In sum, at both extremes of *p*_mix_ the mixed mode results in higher levels of cultural diversity in the population than the “corresponding” pure mode.

Similarly, at both extremes of *p*_mix_ the mixed mode results in a faster rate of cultural change than the “corresponding” pure mode.

### 3.2 Inference

In the previous section we have shown that the different transmission modes result in different patterns at the population level. Here we investigate whether the differences are large enough to ensure distinguishability based on the statistics introduced in Section 2.2.1.

#### 3.2.1 Pure modes

Following the procedure described in Section 2.2.2, we compare the probability distributions of the statistics for pairs of transmission modes *x* and *y,* with *x, y  * {*v, h, o, n*}, and parameter constellations N = 50; μ *=* 0.01; p_w_ = 0.5 and *N* = 50; *μ*; = 0.1; *p*_w_ = 1. Analysis of a larger set of parameters can be found in section S3.3.1 in the supplementary material. Qualitatively, comparison of vertical transmission to the other modes captures the effect of interactions that occur during the individuals’ lifetime. Comparison of age-neutral to oblique and horizontal transmission captures the effect of restricting the set of potential interaction partners to individuals within the same age group and to older individuals, respectively. Finally, comparison of horizontal to oblique transmission captures the effect of the age group of the interaction partner.

We start with analysing the area of overlap *O*_*xy*_ based on the joint probability distribution of the five variants of a trait (Figure 3(a)). Under scenarios with low potential for cultural change (e.g. low *μ*, low *p*_w_; Section 2.1.3), the different transmission modes result in comparable joint probability distributions: *O*_*xy*_ approaches 1 (see Figure S4 in the supplementary material). *O*_*xy*_ tends to decrease as the potential for cultural change increases (i.e. as *μ* and/or *p*_w_ increase). In particular, oblique transmission tends to deviate from the other modes. However, the differences are not large enough to ensure distinguishability based on an arbitrary threshold *Ō* = 0.2 (see the values of *O*_*xy*_ for *f*_joint*, x*_ in Table 3).

**Figure 3:**
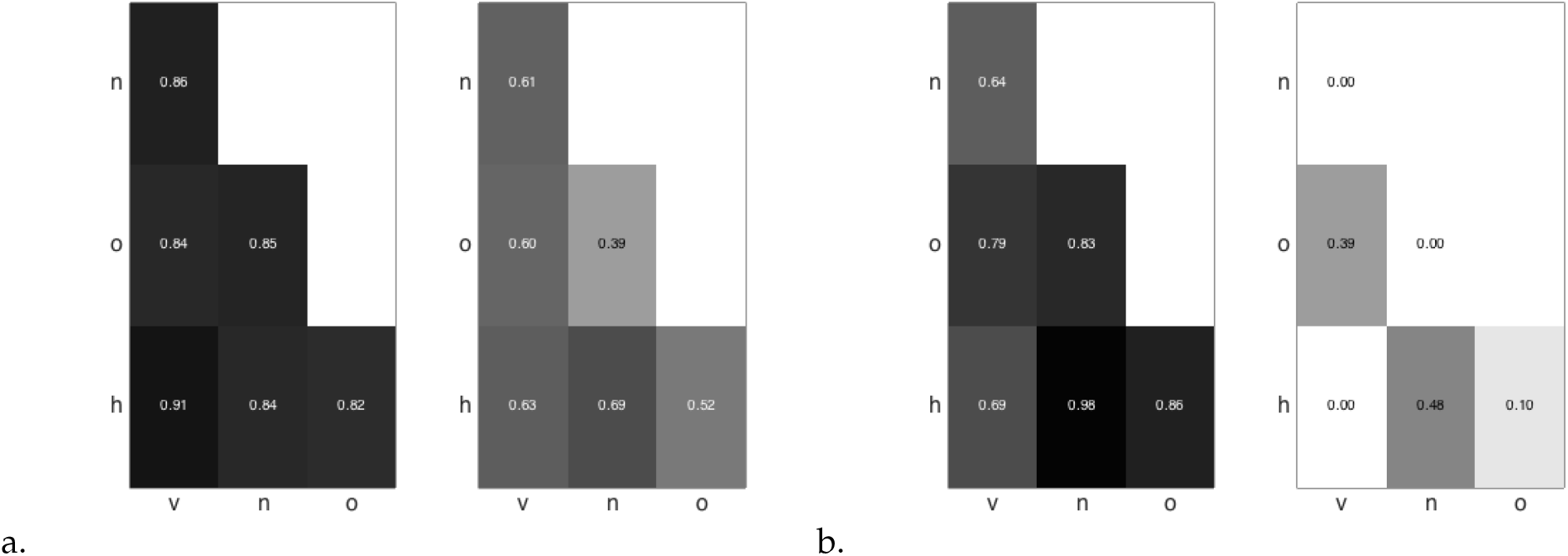
Distinguishability between the pure transmission modes *v, h, o, n* (Table 1) based on (a) the joint probability distribution of the five variants of a trait in the population, and (b) the average time a variant stays the most common variant, 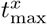. Shown are the values of the area of overlap *O_xy_* between the probability distributions of the statistic for pairs of modes, under parameter constellations *N* = 50; *μ* = 0.01; *p_w_* = 0.5 (left panels) and *N* =50; *μ* = 0.1; *p_w_* = 1 (right panels). The corresponding gray scale ranges from white for *O_xy_* = 0 to black for *O_xy_* = 1.

**Table 3:**
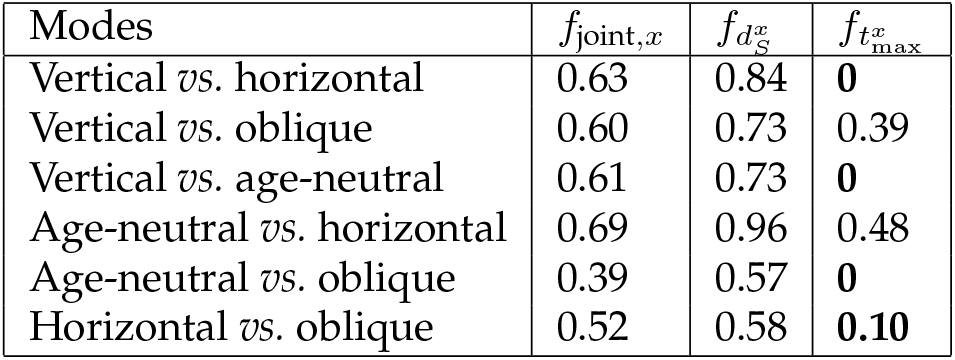
Distinguishability between the pure transmission modes based on select statistics: the joint probability distribution of the five variants of a trait in the population, *f*_joint,*x*_, the level of cultural diversity, *d_S_^x^*, and the average time a variant stays the most common variant, 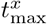. Shown are the values of the area of overlap *O_xy_* between the probability distributions of the statistic for pairs of modes, under parameter constellation *N* = 50, *μ* = 0.1, *p*_w_ = 1. Values in bold indicate distinguishability based on threshold *Ō* = 0.2.

| Modes | $f_{\text{joint},x}$ | $f_{d_S^x}$ | $f_{t_{\max}^x}$ |
| --- | --- | --- | --- |
| Vertical *vs.* horizontal | 0.63 | 0.84 | **0** |
| Vertical *vs.* oblique | 0.60 | 0.73 | 0.39 |
| Vertical *vs.* age-neutral | 0.61 | 0.73 | **0** |
| Age-neutral *vs.* horizontal | 0.69 | 0.96 | 0.48 |
| Age-neutral *vs.* oblique | 0.39 | 0.57 | **0** |
| Horizontal *vs.* oblique | 0.52 | 0.58 | **0.10** |

As noted in Section 2.2.1, the joint probability distribution of the five variants of a trait captures the most information about the system at a given point in time. By contrast, statistics 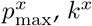 rely on partial data, whereas 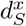 provides summary information. Unsurprisingly, then, the distributions of 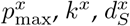 carry a weaker signature of the underlying transmission modes than the joint probability distribution (e.g. compare the values of *O*_*xy*_ for *f*_joint, *x*_ *vs f_d_s_^x^_* in Table 3; the corresponding plots are in Section S3.3.1 in the supplementary material).

Figure 3(b) shows the behaviour of *O*_***xy***_ based on statistic 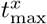, the average time a variant stays the most common variant in the population (Section 2.2.1). Under scenarios with low potential for cultural change (e.g. low *μ,* low *p*_w_), the different transmission modes result in comparable distributions of 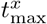 Under scenarios with high potential for cultural change (e.g. high *μ*, high *p*_w_), four pairs (vertical *vs*. horizontal, vertical vs. age-neutral, age-neutral *vs*. oblique, horizonatal *vs*. oblique) produce distributions of 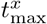 with no overlap (i.e. *O_xy_* = 0; Table 3), indicating that they can be distinguished based on this statistics. In other words, these transmission modes result in temporal dynamics that are sufficiently different to be distinguishable.

Two pairs of modes result in comparable distributions (vertical *vs*. oblique transmission, age-neutral *vs*. horizontal transmission), with areas of overlap substantially greater than 0 (Figure 3(b)). To obtain further insight, we calculate the conditional probabilities *P*(*y*|*z*) given in equation (2) for these pairwise comparisons. Figures 4(a) and (b) show the probabilities based on *t*_max_. In the case of vertical *vs*. oblique transmission (Figure 4(a)), smaller values of empirical estimates of *t*_max_ point to oblique transmission (cf. *P*(*v*|*t*_max_ = *z*) ≈ 0 for small values of *t*_max_), whereas larger values point to vertical transmission (cf. *P*(*v*|*t*_max_ = *z*) ≈ 1 for large values of *t*_max_). We note that in this context “small” and “large” refer only to the comparison between values generated by vertical and oblique transmission. As discussed in Section 3.1, oblique transmission produces larger values of *t*_max_ than horizontal and age-neutral transmission (see Figure S1(d) in the supplementary material). The distinction between age-neutral and horizontal transmission remains ambiguous, however; almost all values of *t_max_* can be produced by either mode (Figure 4(b)). Similar conclusions apply for the level of cultural diversity *d_S_^x^* (Figures 4(c) and (d)).

**Figure 4:**
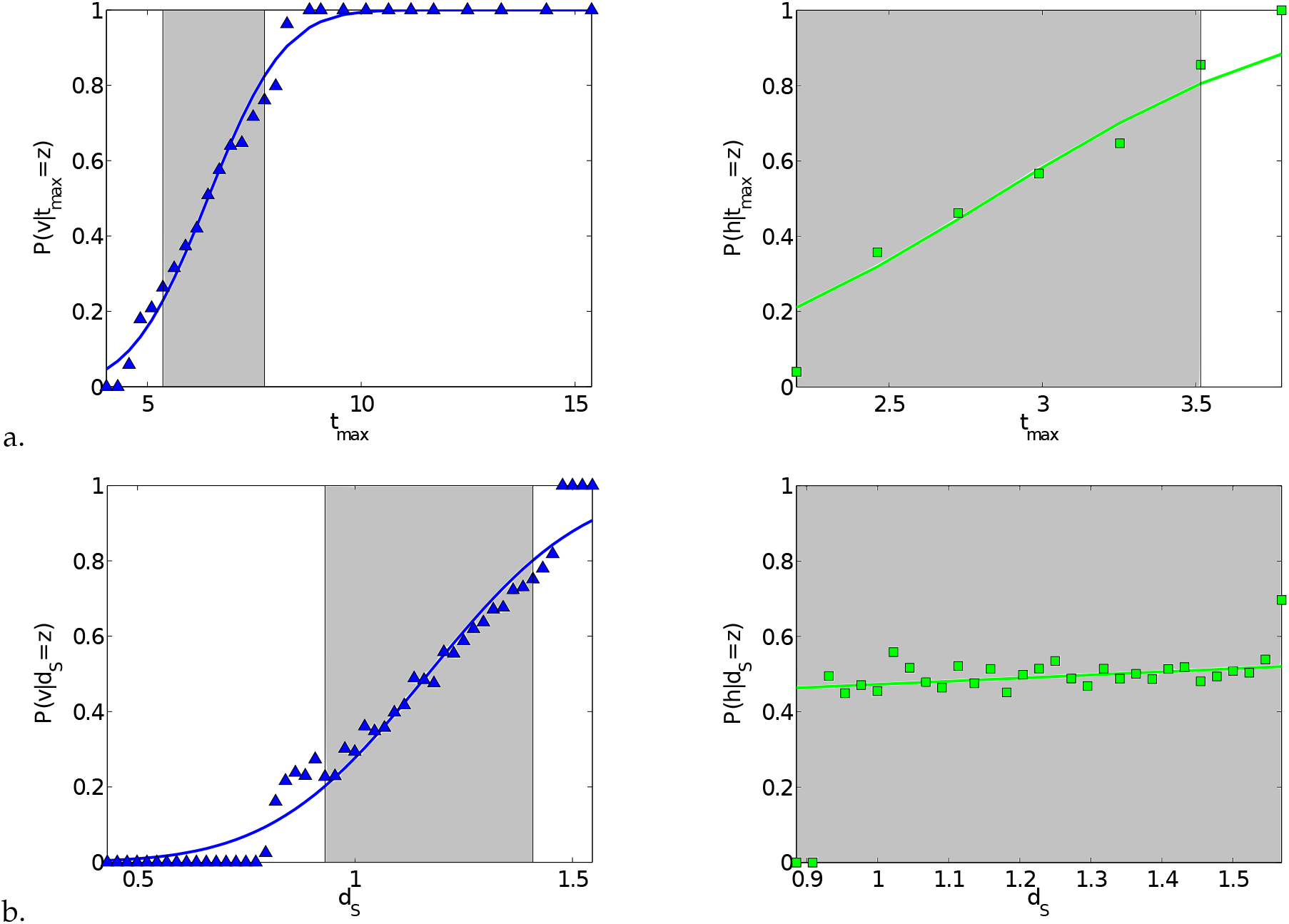
*P*(*x|z*) for pairwise comparisons between vertical and oblique transmission (left panels, blue triangles) and age-neutral and horizontal transmission (right panels, green squares) based on (a) the average time a variant stays the most common variant, 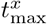, and (b) the level of cultural diversity, *d_S_*, under parameter constellation *N =* 50; *μ=* 0.1; *p_w_ =* 1. The grey areas indicate the values of the statistic which could have been produced by either transmission mode with a probability greater than 0.2.

In sum, the existence of an empirical estimate greatly improves our ability to distinguish between different transmission modes in situations where a *priori* definitions are not sufficient (i.e. in situations where the area of overlap between the corresponding distributions is larger than 0).

#### 3.2.2 Mixed mode

The results for mixed transmission are in Section S3.3.2 in the supplementary material. The aim in this case is to compare distributions of a statistic for (i) the mixed mode at a given level of *p*_mix_ to the pure modes, and (ii) the mixed mode at a given level of *p*_mix_ to the mixed mode at other levels of *p*_mix_.

Consistent with results for the pure modes, the temporal dynamic of cultural change carries more information about the underlying process than the cultural composition of the population at a specific point in time, as captured, respectively, by statistics 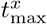 and *d_S_^x^*. Overall, for both statistics the overlap between distributions tends to decrease as the potential for cultural change increases. However, only a limited number of pairs of modes result in distributions that can be distinguished based on the arbitrary threshold *Ō =* 0.2, and only for 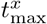. For instance, mixed transmission with *p*_mix_ *=* 0.5 can only be reliably distinguished from vertical transmission. Further, as expected at both extremes of *p*_mix_ the mixed mode cannot be distinguished from the “corresponding” pure mode (i.e. horizontal transmission for low values of *p*_mix_, oblique transmission for high values of *p*_mix_).

#### 3.2.3 Conformity bias

Conformist transmission is known to reduce cultural diversity: the cultural composition of a population tends to be more homogeneous with conformity bias than without it [e.g. 8]. But how strong does the bias have to be to produce characteristic population-level outcomes? In this section we explore whether the presence of conformity bias can be established based on the level of cultural diversity in a population or the temporal dynamic of cultural ange.

Without conformity bias, interactions occur at random between individuals in the relevant age group(s). By contrast, with conformity bias the probability of an individual interacting with a partner carrying a given variant increases with the frequency of the variant in the population (see equation (1)).

Following the procedure described in Section 2.2.2, we compare the probability distributions of statistics 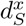 and 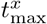 for transmission modes *x, y ∈* {*v, h, o, n*} and parameter constellations *N* = 50, 100; *μ* = 0.1; *p_w_* = 1. Analysis of a larger set of parameters can be found in Section S3.3.3 of the supplementary material. The aim in this case is to compare the distributions of each statistic for a given mode without conformity bias to its conformist counterparts with varying levels of *b* (i.e. *b* = 0 *vs*. *b* = 0.01, 0.02, 0.03; recall that parameter *b* captures the strength of the bias). Note that we focus on scenarios with interaction probability *p*_w_ = 1.

Figure 5(a) shows the areas of overlap *O_xy_* for statistic 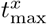, the level of cultural diversity in the population. By definition, vertical transmission is not affected by conformity bias (not shown). For all other modes, as expected there is a trend towards more homogeneous cultural compositions with increasing conformity bias. Thus, as *b* increases, diversity decreases, the distributions of *d_S_^x^* with and without bias become increasingly different, and the values of *O_xy_* decrease as a result. However, the differences are typically not large enough to ensure distinguishability based on an arbitrary threshold *Ō*= 0.2. The only exception is age-neutral transmission for a limited subset of parameter values involving a large population size, low to intermediate mutation rates, and moderate to strong conformity bias (see Figure S6 in the supplementary material). As expected, analysis of *P*(*y*|*z*) given in equation (2) reveals that low values of *d_S_* are consistent with a hypothesis of conformist transmission.

**Figure 5:**
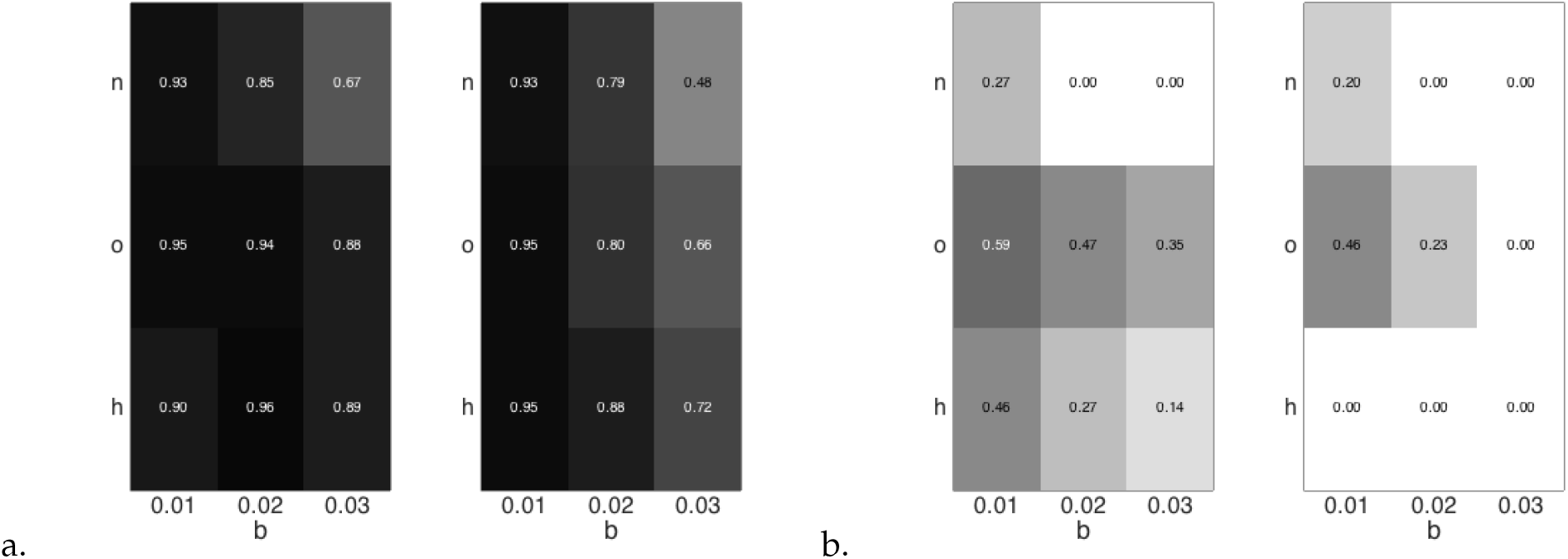
Distinguishability between pure transmission modes *h, o, n* (Table 1) and their conformist counterparts based on (a) the level of cultural diversity, 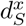, and (b) the average time a variant stays the most common variant, 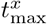. Shown are the values of the area of overlap *O_xy_* between the probability distributions of the statistic for a given mode with and without conformity, with varying levels of *b =* 0.01; 0.02; 0.03, under parameter constellations *N* = 50; *μ =* 0.1; *p_w_* = 1 (left panels) and *N =* 100; *μ =* 0.1; *p_w_* = 1 (right panels). The corresponding gray scale ranges from white for *O_xy_* = 0 to black for *O_xy_* = 1.

Figure 5(b) shows the behaviour of *O_x_*_y_ for statistic 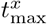 the average time a variant stays the most common variant in the population. We expect this to increases with conformity bias, which by definition “sustains” the most common variant in the population. Our results show that the ability to reliably detect the bias increases with population size. At the same time, it is contingent on the number of potential interaction partners. For example, for age-neutral transmission, moderate conformity can be detected for almost all parameter settings (see also Figure S6 in the supplementary material). By contrast, restricting the set of potential partners to peers (horizontal transmission) or older individuals (oblique transmission) reduces the range of parameter values for which the bias can be detected. In the case of oblique transmission, in fact, only strong bias can be reliably detected, and only for intermediate to high mutation rates (specifically, *N* = 100; *μ* = 0.1; *b* = 0.03). Analysis of *P*(*y*|*z*) reveals that large values of *t*_max_ point to the existence of a conformist bias in the population.

#### 3.2.4 Continuous traits

In section S3.3.4 of the supplementary material we apply the framework described above to continuous cultural traits, i.e. traits that can take any value in a given interval. Overall, we find that the population average of the trait does not carry a detectable signature of the underlying transmission modes. Binning the interval of trait values into discrete variants results in greater inferential power, but the binning must be sufficiently fine-grained, with the traits discretized in more than three variants (see Section S3.3.4 for a detailed analysis).

## 4 Discussion

We developed a simulation model to explore the interface between individual-level processes and population-level patterns in human cultural evolution. Our first aim was to derive broad expectations about different pathways for the transmission of information between individuals. Our second aim was to establish theoretical limits to inference based on the population-level patterns produced by these pathways. Are the patterns different enough that they can be reliably distinguished based on population-level statistics, providing insight into the underlying individual-level processes?

The model tracks the distribution of variants of cultural traits across individuals in a population over time, conditioned on different modes for the transmission of information between individuals, for a range of parameters capturing demographic and cultural factors. Building on previous work [7], we investigated four “pure” modes: vertical (parent to offspring), horizontal (peer to peer, by age group), oblique (older to younger, excluding parent to offspring), and age-neutral (any individual in the population). A fifth “mixed” mode effectively combined horizontal and oblique transmission. We also investigated the effect of conformity bias, a preference for the most common variant in the population [8].

We used four statistics to summarize the cultural composition of the population at a specific point in time: the frequency distribution of the different variants of the cultural trait, the frequency of the most common variant, the total number of variants present, and the level of cultural diversity. An additional statistic measured the rate of change of the most common variant of a trait. This gave an indication of how fast the cultural composition of the population can change, providing insight into the temporal dynamic of the process.

### 4.1 Age structure, trickle-down effect, and relative rates of change

The five transmission modes differ in the set of potential interaction partners (Section 2.1.1). Previous work suggests that they should therefore produce substantially different evolutionary dynamics, modulated by demographic and cultural factors [e.g. 7]. For example, [18] explored the effects of vertical, oblique, and horizontal transmission on the age-structure of a population, and the extent to which cultural traits affecting demographic change can spread. They showed that a trait that reduces fertility but increases survival can spread to fixation and lead to substantial demographic change, in the form of an increase in population size, under certain modes of cultural transmission.

Our results, based on analysis of neutral traits and constant population sizes, show that the five transmission modes lead to differences in evolutionary dynamics. These differences are most pronounced under scenarios with high potential for cultural change (e.g. high mutation rates, high interaction probabilities). In particular, we have shown that oblique transmission is strongly affected by the value of the interaction probability: when this is high, oblique transmission results in more homogenous cultural compositions compared to the other pure modes. Broadly, a homogenous cultural composition is reflected in an uneven frequency distribution of the cultural variants (i.e. a low number of variants present due to the high frequency of the most common variant), resulting in low cultural diversity (Section 3.1).

Further, we have shown that this effect is rooted in the interplay between the different sets of potential interaction partners and the age structure of the population. In particular, oblique transmission is characterised by a disproportional influence of older individuals, whereby a variant at high frequency in one age group trickles down to all younger age groups; over time, this leads to a more homogeneous population. For example, an individual in the oldest age group is a potential interaction partner for all younger individuals in the population, whereas a “middle-aged” individual is a potential partner for only the younger half of the population. This effect leaves a signature at the population level in terms of (i) lower levels of cultural diversity, and (ii) slower rates of cultural change, for oblique transmission compared to horizontal and age-neutral transmission. Horizontal and age-neutral transmission show comparable rates of change.

Vertical transmission leads to the slowest rates of change. In our framework, this mode involves a “one-shot” interaction event between parent and offspring, with opportunities for change limited to the process of mutation. The relative “conservativeness” of vertical transmission is a well-established result in the theory of gene-culture coevolution, starting with seminal work by [7]. We return to this issue below, after outlining our results regarding inference.

### 4.2 Inferring process from pattern

#### 4.2.1 Summary of results

Questions relating to the indentifiability of transmission modes or other underlying processes are of course not restricted to the field of cultural evolution. For example, a large body of work in population genetics focuses on testing for the presence or absence of selective forces [e.g. 26, 27, 28]. The general idea, similar to the one used here, is to derive expectations for different quantities of interest under a specific evolutionary scenario, and to subsequently compare empirical estimates of the quantities with those expectations. Recently, inferential frameworks have been applied directly to observed data. These frameworks often combine generative modeling of the system under consideration and Bayesian inference techniques. For example, in this way researchers have gained important insights into the transmission dynamics of infectious diseases and other epidemiological processes [e.g. 29, 30, 31], and into human demographic history [e.g. 32, 33, 34]. Rather than focusing on a particular cultural dataset [see e.g. 11, 12, 35, 36, 37, for examples of such analyses], our aim here was to develop a theoretical understanding of which transmission modes can be distinguished on the basis of population-level data. This relates to the problem of equifinality, i.e. which modes can produce similar population-level patterns.

Our analysis provides two kinds of results: first, general expectations about the distinguishability of population-level outcomes produced by the different transmission modes in the absence of empirical data; second, given an empirical estimate of a statistic, the probability that a given transmission mode acted in the population to produce this estimate.

Beginning with the pure modes, our results show that under scenarios with low potential for cultural change (e.g. low mutation rates, low interaction probabilities) the modes produce outcomes that cannot be reliably distinguished based on any statistic. Yet outcomes tend to diverge as the potential for cultural change increases. In particular, under scenarios with high potential for cultural change they can be reliably distinguished, in most cases, based on the temporal dynamic. The two exceptions are vertical *vs*. oblique transmission, and age-neutral vs. horizontal transmission. In these two cases, additional insight can be gained from empirical estimates of the average time a variant stays the most common variant. For vertical *vs*. oblique transmission, small values are indicative of oblique transmission, whereas large values clearly point to oblique transmission. For age-neutral vs. horizontal transmission, even an empirical estimate will likely not be able to resolve the distinguishability issue.

These results suggest that even when outcomes are similar in terms of cultural compositions, they can differ substantially in temporal dynamics — in other words, similar distributions of cultural variants at a specific point in time can be reached through substantially different processes. In sum, for the pure modes the temporal dynamic of cultural change over time retains a stronger signature of the underlying processes than a “snapshot” of the relative frequencies of the variants at a given point in time, and the signature is stronger the greater the potential for cultural change.

This general insight also applies to mixed transmission. However, in this case even under scenarios with high potential for cultural change the signature retained by the temporal dynamic tends to be too weak for outcomes to be reliably distinguished.

Finally, we have shown that the signature of conformity bias is in general stronger for the temporal dynamic than for the level of cultural diversity. As expected, the level of cultural diversity decreases with increasing conformity bias, while the temporal dynamic slows down. At the same time, with small population sizes the signature of conformist transmission tends to be obscured by random drift. It becomes stronger as population size increases, especially for age-neutral transmission, as the resulting increase in the pool of potential interaction partners is larger in this case than for horizontal or oblique transmission. For horizontal and oblique transmission, only strong conformity bias can be reliably detected, in most cases, and only based on the temporal dynamic with large population sizes.

Empirical estimates can provide further insight also in this case. Specifically, low values of cultural diversity and large average times a variant stays the most common point strongly to the existence of a conformist bias, when compared with the non-conformist situation.

#### 4.2.2 Implications for empirical studies

We conclude by reviewing the implications of our findings for empirical studies of cultural evolution. Broadly, our work complements alternative theoretical approaches used to infer process from pattern in human culture, including efforts based on adoption curves [e.g. 16, 38, 39, 40], rank-abundance distributions [e.g. 41, 19, 42], levels of diversity [e.g. 43, 1, 44], and turnover rates [e.g. 45, 46], as well as model selection frameworks [e.g. 47].

In terms of inference, our study suggests that if the frequency distribution of the different variants of a trait is available, then inference procedures should rely on these data where possible. Unsurprisingly, using partial data (e.g. the frequency of the most common variant of a trait) or summary statistics (i.e. the level of cultural diversity) results in loss of useful information. For example, a diversity measure such as the Shannon index is often used to summarise the cultural composition of a population [e.g. 1, 23, 24]. We have shown that the dimension reduction involved in its calculation obscures population-level differences in the patterns produced by the transmission modes.

As discussed above, our results suggest that the temporal dynamic of cultural change over time is more instructive about the underlying processes than the cultural composition of the population at a given point in time. The approach we used to characterize the temporal dynamic is more parsimonious than other approaches in the literature. For example, [45] use the turnover rate, defined as the number of new variants of a trait that enter the list of variants at the highest frequency in each time step (analogous to the “new entries” in a top-10 or top-100 chart). However, this measure does not readily extend to traits that include only a small number of variants (e.g. five variants, as in our case). Additionally, it requires information about the frequencies of all variants in the population. By contrast, our approach only requires information about the frequency of the most common variant.

At the same time, it should be noted that the results presented here rest on the assumption that we have complete information about the most common variant of the trait at every point in a given time interval — in other words, that estimation of the temporal dynamic of cultural change is exact. Such detailed time series data are difficult to obtain, however, especially for existing datasets and/or those that rely on historical information. In a related study using the mathematical framework developed here, we investigated how sparse the time series data can be for the transmission modes to still be distinguishable [48]. Results show that if only incomplete information is available, i.e. the most common variant is known for a sample of time points, then the level of distinguishability depends on the properties of the sample. In particular, the distance between the time points affects how much insight can be obtained from population-level data.

In general, if information about the most common variant in the population is only available for a sample of time points, one should not expect levels of distinguishability comparable to those we report here.

In practice, the use of the most appropriate statistic or inference framework is dictated by the available data. Given our focus on cultural evolution, we have analyzed statistics that can potentially be derived even from sparse data — a common feature of datasets in athropology and related disciplines. Where researchers have access to data with high temporal resolution, such as time series describing the frequency change of different cultural variants, then adoption curve analyses, and especially model selection frameworks, can be instructive.

A final set of insights relates to the analysis of continuous cultural traits. Our results show that the mean value of a trait across the population does not carry a detectable signature of the underlying transmission modes; therefore, it is not a useful statistic for characterising the cultural composition of a population. Binning the interval of trait values into discrete variants results in greater inferential power, but the binning must be sufficiently fine-grained. For example, discretisation into two or three variants (corresponding to e.g. present *vs.* absent or small *vs.* medium *vs.* large) is generally not enough to ensure distinguishability based on the temporal dynamic of cultural change, whereas discretization into five or ten variants produces distinguishability results comparable to the discrete case.

We conclude with a general observation, bearing on the interface between the observed population-level patterns and the underlying individual-level processes. As discussed above, our results show that vertical transmission leads to the slowest rate of change of all the modes, consistent with the notion prevalent in the literature of its relative “conservativeness” [7]. This notion continues to provide the foundation to a large body of empirical work, including field-based investigations [e.g. 10, 11, 13] and cross-cultural studies [e.g. 12, 14, 15, 12]. At the same time, our inference results show that vertical transmission can produce temporal dynamics similar to other modes, and in particular to oblique transmission. Empirical estimates of statistics describing the temporal dynamic can provide further crucial information. Still, our findings invite caution in linking population-level patterns to individual-level processes based on data documenting variation in cultural traits within and between populations.

This example illustrates well the theoretical limits to inferring individual-level processes from population-level patterns in human cultural evolution. In particular, we should not expect a one-to-one mapping between population-level statistics and the underlying transmission modes: different scenarios can lead to comparable patterns at the level of groups. Consistency between any one specific scenario and empirical data should be interpreted in this context. However, acknowledging the problem of equifinality does not imply that we cannot extract any information about cultural evolution from these data. Mathematical frameworks similar to one used here can provide general expectations, in form of probability distributions, against which to compare empirical estimates. Further, statistical inference procedures that compare simulated data to empirical data can help delimit the amount of information that can be extracted on a case-by-case basis [e.g. 35, 36, 37]. Conceptually, this shifts the focus from identifying the one scenario that likely produced the observed data to excluding those that likely did not.

## Acknowledgement

This work was supported by a National Science Foundation Early Concept Grant for Exploratory Research to LF and AK (EAGER 1249146, “Linking Pattern and Process in Cultural Evolution”). We thank Tanmoy Bhattacharya and Mark Galassi for insightful discussions and Adam Powell for helpful comments on an earlier version of this manuscript.

